# Ventromedial prefrontal cortex shields value computations from bodily arousal in decision making

**DOI:** 10.1101/2025.11.07.687164

**Authors:** Patricia Christian, Jakob Kaiser, Georgia Eleni Kapetaniou, Simone Schütz-Bosbach, Alexander Soutschek

## Abstract

Emotional reactions to norm violations influence how people trade-off selfish interests against social norms, but it remains unknown which brain regions causally contribute to integrating physiological arousal associated with norm violations into the decision process. Here, we provide evidence that the ventromedial prefrontal cortex (vmPFC) causally contributes to decoupling value computations in normative choice from physiological arousal. Excitatory brain stimulation over the vmPFC reduced aversion to disadvantageous inequality in an Ultimatum game and increased post-decision confidence after acceptance of unfair monetary splits. Physiological arousal as measured via heart rate changes was increased in response to splits perceived as unfair. Enhancing vmPFC excitability reduced the sensitivity to unfairness not by directly inhibiting physiological arousal but by blocking its influence on the decision process. This suggests that vmPFC causally moderates how physiological responses to norm violations affect trade-offs between self-interests and norm considerations, highlighting its key role for integrating emotional and cognitive processes in decision making.

**Significance statement:** Emotions like anger about the violation of social norms are associated with physiological arousal, and the interoceptive awareness of this arousal is thought to influence decision making. Here, we identify the neural mechanism that causally mediates the influence of physiological arousal on the decision process. Increasing the excitability of the ventromedial prefrontal cortex strengthened selfish interests over fairness considerations in social interactions via decoupling the decision process from unfairness-evoked arousal. This provides a neuroscientific explanation for how physiological arousal influences social decisions and suggests that the role of the ventromedial prefrontal cortex for the interaction between emotion and cognition in decision making is based on interoception rather than emotion regulation.

## Introduction

People often have to trade-off their selfish interests against fairness considerations (Fehr & Camerer, 2007; Guth, Schmittberger, & Schwarze, 1982). For example, if a person treats us unfairly, we may punish the other for her norm violation even at our own expenses. Experiencing norm violation is often associated with enhanced physiological arousal that accompanies emotions like anger about the unfair treatment; this physiological reaction is thought to affect the value we assign to punishing the norm violation (Dunn, Evans, Makarova, White, & Clark, 2012). Little is known, however, about the neural mechanisms that moderate the influence of emotion-related physiological arousal on decisions involving trade-offs between selfish interests and fairness considerations.

The ventromedial prefrontal cortex (vmPFC) fulfills the criteria required to play a central role in integrating physiological arousal into the decision process. Theoretical accounts ascribe the vmPFC important contributions to emotion processing (Hiser & Koenigs, 2018; Rudebeck, Bannerman, & Rushworth, 2008), according to which the vmPFC may moderate the influence of emotion-related physiological arousal on decisions in different ways: First, the vmPFC might directly be involved in emotion regulation, as suggested by evidence linking the vmPFC to the voluntary down-regulation of negative emotions (Egner, Etkin, Gale, & Hirsch, 2008; Etkin, Buchel, & Gross, 2015; Suzuki & Tanaka, 2021). In keeping with this, lesions to vmPFC increased physiological responses to emotionally salient stimuli (Jenkins et al., 2018). From this perspective, vmPFC may attenuate the impact of norm violations on choice behavior by inhibiting the physiological response to perceived unfairness.

Alternative accounts posit the vmPFC to be involved in interoception rather than emotion regulation, that is, the subjective awareness of physiological responses like heartbeat changes (Azzalini, Buot, Palminteri, & Tallon-Baudry, 2021; Babo-Rebelo, Richter, & Tallon-Baudry, 2016; Park, Correia, Ducorps, & Tallon-Baudry, 2014). Interoception was found to increase the influence of unfairness-elicited bodily responses on social decisions (Dunn et al., 2012), though conflicting evidence suggests that interoception may alternatively also reduce the influence of emotional reactions to unfairness on decision making (Kirk, Downar, & Montague, 2011). This account predicts that vmPFC activity does not alter the physiological response to unfairness itself but moderates (either strengthens or weakens) its impact on decisions to punish unfair behavior. Thus, while emotion regulation and interoception were claimed to be at least partially intertwined constructs because interception can improve emotion regulation (Critchley & Garfinkel, 2017), these accounts make nevertheless empirically distinguishable predictions: if vmPFC activation influences emotion regulation processes, experimentally manipulating vmPFC activation should change unfairness-related physiological arousal. In contrast, if vmPFC implements interoceptive but not emotion regulation processes, vmPFC activation should not alter the physiological response per se but moderate its influence on decision making.

Existing evidence on the role of the vmPFC for trading-off fairness versus selfish interests provided no conclusive support for one of these alternatives. While one line of evidence linked the vmPFC to a stronger weighting of selfish interests over fairness norms (Gilam et al., 2017; Koenigs & Tranel, 2007), other studies suggest the vmPFC to increase the sensitivity to norm violations (Baumgartner, Knoch, Hotz, Eisenegger, & Fehr, 2011; Feng, Luo, & Krueger, 2015; Gu et al., 2015; Lo Gerfo et al., 2019). However, the inferences that can be drawn from these findings remain limited because they either did not causally test the contribution of the vmPFC (Baumgartner et al., 2011; Feng et al., 2015) or they lack the necessary spatial resolution to conclusively link changes in social decision making to vmPFC activation (Gilam et al., 2017; Gu et al., 2015; Koenigs & Tranel, 2007; Lo Gerfo et al., 2019). Moreover, no previous study assessed physiological responses to norm violations, as would be necessary to decide whether the vmPFC affects decision making via down-regulating physiological arousal (emotion regulation account) or via moderating its influence on the decision process (interoception account). Thus, it remains unknown whether, and if so how, the vmPFC causally contributes to integrating unfairness-induced physiological responses into trade-offs between selfish interests and fairness considerations.

The goal of the current study was to decide between these competing accounts of vmPFC involvement in normative decision making. To determine how the vmPFC moderates the influence of physiological arousal on normative decision making, participants played an Ultimatum game after either anodal or sham transcranial direct current stimulation (tDCS) targeting the vmPFC. Participants had to decide whether to accept or reject unfair monetary splits of anonymous human proposers. We measured changes in heart rate during the presentation of the offers as indicator of the physiological response to unfairness. Previous studies found that increases in heart rate are associated with rejections of unfair offers, supporting the view that they reflect physiological arousal associated with the negative emotional response to inequality (Gilam et al., 2017; Sarlo, Lotto, Palomba, Scozzari, & Rumiati, 2013). Crucially, measuring heart rate responses as physiological reaction to unfairness allowed us to decide between different functional roles of the vmPFC: If the vmPFC down-regulates emotional responses to unfairness, we would expect anodal tDCS over vmPFC to reduce unfairness-related changes in heart rate. Alternatively, if the vmPFC moderates the influence of arousal on the decision process via interoceptive processes, enhancing vmPFC excitability should change the influence of physiological arousal on decisions in the Ultimatum game. Moreover, given the vmPFC’s involvement in representing subjective post-decision confidence (Vaccaro & Fleming, 2018), we also hypothesized that vmPFC tDCS strengthens post-decision confidence via lowering the sensitivity to fairness violations.

## Materials and Methods

### Participants

Thirty-five healthy volunteers (mean age = 26.5 years, range years = 18-35, 16 female, 19 male) who were recruited at the Ludwig Maximilians University (LMU) Munich participated in the study. We excluded data from 3 participants due to technical issues with the heart rate recordings, leaving 32 participants for the analysis. According to a power analysis, 20 participants are sufficient to detect a significant effect (alpha = 5%, power = 95%) assuming an effect size of Cohen’s d = 0.86 observed in a meta-analysis of brain stimulation studies in social decision making (Christian & Soutschek, 2022). This sample size provides also sufficient power according to a previous tDCS study on the Ultimatum game (Gilam et al., 2017). All participants were healthy volunteers without any known psychiatric or neurological disorders and were naive with respect to the aims of the study. Ethical approval was granted by the ethics committee of the psychology department at LMU and all procedures followed the principles of the Declaration of Helsinki (2013). Participants provided informed written consent prior to participation. Participants showed no contra-indications for tDCS and were informed about its potential side-effects (Bikson et al., 2016). As compensation, participants received 10 euro per hour and a monetary bonus depending on their choices (see below).

### Stimuli and task design

The study followed a randomized, sham-controlled, single-blind crossover design. All participants took part in two experimental sessions where they received either anodal or sham tDCS over vmPFC (in counterbalanced order) prior to performing the Ultimatum game (Figure 1A). Participants played the Ultimatum game in the role of the responder and had to decide whether to accept or reject a split of 20 euro offered by an anonymous human proposer. The amount offered ranged from 0 (unfair offer) to 10 euro (fair offer) in steps of 1 euro. If participants accepted an offer, the participant and the proposer obtained the indicated amounts of money; if an offer was rejected, none of them obtained money. Participants therefore had to trade-off their selfish interests (accepting an offer to obtain the bonus) against the costly punishment of norm violations (rejection of unfair offers). It is important to emphasize that choices had real consequences for participants’ and proposers’ payments. In a pre-study, we had collected offers from various proposers, which we then used for the Ultimatum game in the main study. We informed participants that the offers were made by real individuals in pre-study and that, at the end of the experiment, one randomly selected decision would be paid out to the participant and a proposer from the pre-study.

**Figure 1.**
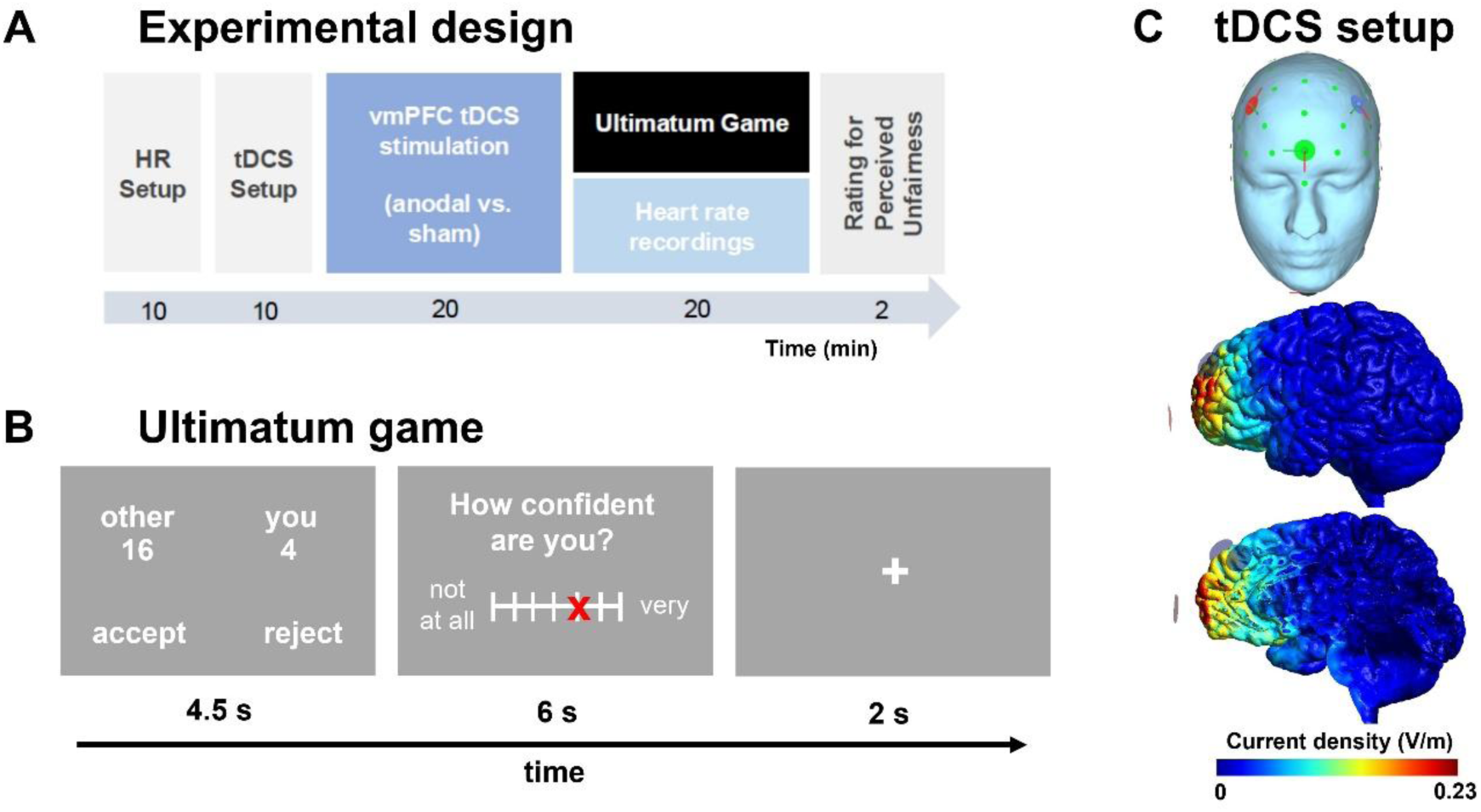
(A) Experimental procedure. Participants completed two separate sessions, in which either anodal or sham offline tDCS was applied over the ventromedial prefrontal cortex (vmPFC) for 20 minutes in a counterbalanced within-subject design. After stimulation, participants provided a brief rating of tDCS-induced perceived discomfort and flickering. Following this, participants performed the Ultimatum Game for 20 minutes while their heart rate was continuously recorded. Upon completing the task, participants indicated their perceived unfairness of the offers made by the proposers. (B) Example trial for the Ultimatum Game. Participants were presented with proposals regarding how to split an amount of 20 euros, made by an anonymous proposer. Participants had to decide whether to accept or reject the offer. After each choice, participants rated their subjective confidence to have made the best choice. (C) tDCS setup: We targeted the vmPFC with a high-definition electrode setup. Current flow simulations with the SimNIBS toolbox (Saturnino et al., 2019) suggest that current density was highest (indicated by warmer colors) in ventral parts of the medial prefrontal cortex.

On each trial, we first presented participants an offer (e.g., “4 euro for you, 16 euro for other”) on a screen. Participants had to indicate their choice to accept or reject the offer by pressing the left or right arrow key (key-response assignment was counterbalanced across participants) within 4.5 seconds. Afterwards, participants had to rate their confidence in having made the best choice on a continuous Likert scale ranging from 0 (not confident at all) to 100 (very confident) within 6 seconds, followed by an inter-trial interval of 2 seconds (Figure 1B).

### Procedure

All participants took part in two experimental sessions in which they received either anodal or sham tDCS over vmPFC (in a counterbalanced order). Offline tDCS was applied for 20 minutes prior to the Ultimatum Game (see below for details). Following tDCS, participants performed 96 trials of the Ultimatum Game (approx. 20 minutes) in randomized order while we continuously recorded their heart rate. At the end of the experiment, participants were asked to rate as how unfair they perceived each offer on a continuous scale from 0 (not unfair at all) to 100 (very unfair). These ratings were collected after the Ultimatum game to ensure that they did not influence participants’ choices in the Ultimatum game. At the end of the experiment, one choice was randomly selected for the bonus payment.

### tDCS protocol

We used a 16-channel tDCS stimulator (neuroConn, Ilmenau, Germany) to apply tDCS with a current strength of 1.5 mA (peak-to-peak) for 20 minutes in line with the safety recommendations for tDCS (Bikson et al., 2016; Woods et al., 2016). This protocol is thought to result in tDCS-induced aftereffects for at least 30 minutes (Kuo et al., 2013; Polanía, Nitsche, & Ruff, 2018). We placed one round high-definition anode (diameter = 2 cm) over the vmPFC (position Fpz according to the international EEG 10/20 system), surrounded by 3 cathodes over F3, F4, and under the chin. The electrode under the chin allowed us to induce a current flow deeper into the brain to stimulate large parts of vmPFC (Kroker et al., 2022; Martin, Huang, Hunold, & Meinzer, 2019) (Figure 1C). To verify that tDCS targeted the vmPFC, we performed a region of interest (ROI) analysis of the electric fields generated by tDCS using the SimNIBS toolbox (Saturnino et al., 2019). The vmPFC ROI (MNI coordinates: x = -3, y = 54, y = -12) was defined based on a previous study reporting the vmPFC to correlate with offer fairness in the Ultimatum game (Baumgartner et al., 2011). The electric field in the vmPFC ROI was 0.13 V/m, whereas the field strength in an ROI for the right dlPFC (x = 45, y = 24, z = 21) from the same study was only 0.06 V/m. This suggests our tDCS setup to show pronounced effects on vmPFC, but not dlPFC.

For anodal tDCS, the stimulation lasted a total of 20 minutes (ramp-up: 15 seconds, ramp-down: 15 seconds). Note that we employed anodal rather than cathodal tDCS because cathodal tDCS was reported to often result in paradoxical excitatory effects (Jacobson, Koslowsky, & Lavidor, 2012). In the sham condition, the current was ramped up for 15 seconds and directly afterwards ramped down for 15 seconds. In the following 20 minutes, the sham protocol followed a repeated pattern of stimulation-free intervals of 28 seconds, ramp-up phases of 1 second, directly followed by ramp-down phases of 1 second. This was done to improve participants’ blinding to the current tDCS condition. At the end of the stimulation, participants had to rate the perceived discomfort and flickering during the stimulation on rating scales from 0 (not at all) to 7 (very much) (time limit = 4 seconds). We observed no significant differences in perceived discomfort, *t*(31) = 1.80, *p* = 0.08, or flickering, *t*(31) = 1.83, *p* = 0.08, between anodal and sham tDCS. Nevertheless, we entered discomfort and flickering ratings as control variables to all statistical analyses to rule out that any influences of tDCS on choice behavior might be related to the perceived discomfort of the stimulation.

### ECG protocol

Continuous electrocardiography (ECG) data were recorded with two bipolar electrodes below the left pectoral muscle and the left clavicle, as well as one ground electrode below the right clavicle. ECG data was recorded with a Brain-Vision QuickAmp amplifier, using a sampling rate of 500 Hz.

### ECG analysis

Preprocessing of ECG data was performed using the FieldTrip toolbox (Oostenveld, Fries, Maris, & Schoffelen, 2011) and MATLAB 2021b (MathWorks Inc. Natick, MA, USA). ECG data were filtered using a 40 Hz low-pass filter and a 1 Hz high-pass filter. We extracted heart rate responses from the time window of interest (presentation of offer for 4 seconds). We used the Pan-Tompkins algorithm in the BioSigKit toolbox (Sedghamiz, 2018) for detecting R-peaks in the time window for each trial. Based on these, we calculated the latency between R-peaks within the time window of interest (Kaiser, Belenya, Chung, Gentsch, & Schutz-Bosbach, 2021). We converted the mean latencies between peaks for each trial to beats per minute. Based on the identified R-peaks, we excluded outlier trials for which the heart rate deviated more than 3 standard deviations from participant’s mean (2.9% of all trials).

### Statistical analysis

We analyzed choices, confidence ratings, and heart rate responses in the Ultimatum game with general(ized) linear mixed models (GLMMs) using the lme4 package in R (Bates, Mächler, Bolker, & Walker, 2014). The alpha threshold was set to 5% (two-tailed) for all analyses. To test the hypothesis that vmPFC tDCS changes the acceptance of unfair offers, we regressed binary choices to accept or reject an offer (0 = reject, 1 = accept) on fixed-effect predictors for tDCS (0 = sham, 1 = anodal), Offer (z-transformed amount of money offered to participant), and the interaction term. All predictors were also modelled as random slopes in addition to participant-specific random intercepts. As covariates of no-interest, we controlled for tDCS-induced discomfort and flickering in all statistical analyses. In further GLMMs, the perceived unfairness of offers, heart rate responses, and post-decision confidence were regressed on predictors for tDCS, Offer, Choice (0 = reject, 1 = accept), and all interaction terms. Lastly, to test whether vmPFC tDCS changes the influence of heart rate responses on choices, we conducted a further GLMM where binary choices were regressed on predictors for tDCS, trial-specific Heart rate, Offer, and all interaction terms. This allowed testing the hypothesis that vmPFC moderates the influence of the physiological response to unfairness on decisions in the Ultimatum game.

To further assess whether vmPFC tDCS affects preferences for unfair outcomes, we also fitted an inequality aversion model to the binary choice data with a hierarchical Bayesian approach. In more detail, we assumed that the utility of accepting versus rejecting an offer was given by the following equation:

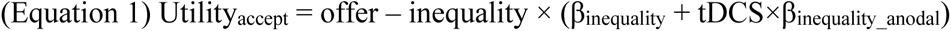

Here, β_inequality_ is a free parameter quantifying the aversion against disadvantageous inequality, tDCS is a dummy-coded variable that is 0 for sham and 1 for anodal tDCS, and β_inequality_anodal_ captures the influence of anodal tDCS on inequality aversion. A preference for accepting or rejecting an unfair offer is indicated by Utility_accept_ > 0 and Utility_accept_ < 0, respectively. We used a standard softmax function to translate utilities into binary choices:

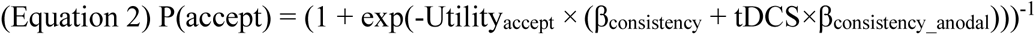

Where β_consistency_ is a choice consistency (“inverse temperature”) parameter indicating how consistently participants chose the higher utility option, whereas β_consistency_anodal_ quantifies the influence of anodal tDCS on choice consistency. For all model parameters, we estimated the parameter values under sham as well as the influences of anodal tDCS. We estimated parameters with a hierarchical Bayesian approach using the dbern module in JAGS. We used non-informative flat priors for the group parameters under sham and normally distributed priors for the tDCS effects. We assumed that individual parameters are normally distributed around the group-level parameters. We computed four chains, each with 30,000 samples (burn-in = 5,000). The *R* values for all parameters were ≤ 1.01, indicating model convergence. For statistical inference, we assessed whether 95% highest density intervals (HDI_95%_) included zero.

## Results

### vmPFC tDCS strengthens self-interests over fairness concerns

First, we aimed to disentangle between conflicting accounts on the role of the vmPFC for normative choice (Baumgartner et al., 2011; Gilam et al., 2017; Koenigs & Tranel, 2007; Lo Gerfo et al., 2019) by testing how anodal vmPFC tDCS influences the acceptance of unfair offers. As to be expected, participants were more willing to accept unfair splits the more money was offered to them, beta = 5.61, *z* = 10.52, *p* < 0.001. Anodal versus sham tDCS over vmPFC increased acceptance rates, beta = 1.40, *z* = 2.84, *p* = 0.005, independently of the fairness of the offer, tDCS × Offer: beta = -0.49, *z* = 0.94, *p* = 0.35 (Figure 2A). Thus, participants’ choices under vmPFC tDCS were more strongly guided by their economic self-interests than by fairness considerations. However, vmPFC tDCS did not affect the perceived unfairness of offers (measured after the decision task but still under the influence of tDCS): While lower monetary offers were considered as increasingly unfair, beta = -26.06, *z* = 22.34, *p* < 0.001, there were no significant tDCS effects on perceived unfairness, beta = -2.86, *z* = 1.82, *p* = 0.08 (Figure 2B). This provides no evidence that enhancing vmPFC excitability alters the normative evaluation of unfair offers per se; instead, vmPFC stimulation changes how fairness is traded-off against selfish interests.

**Figure 2.**
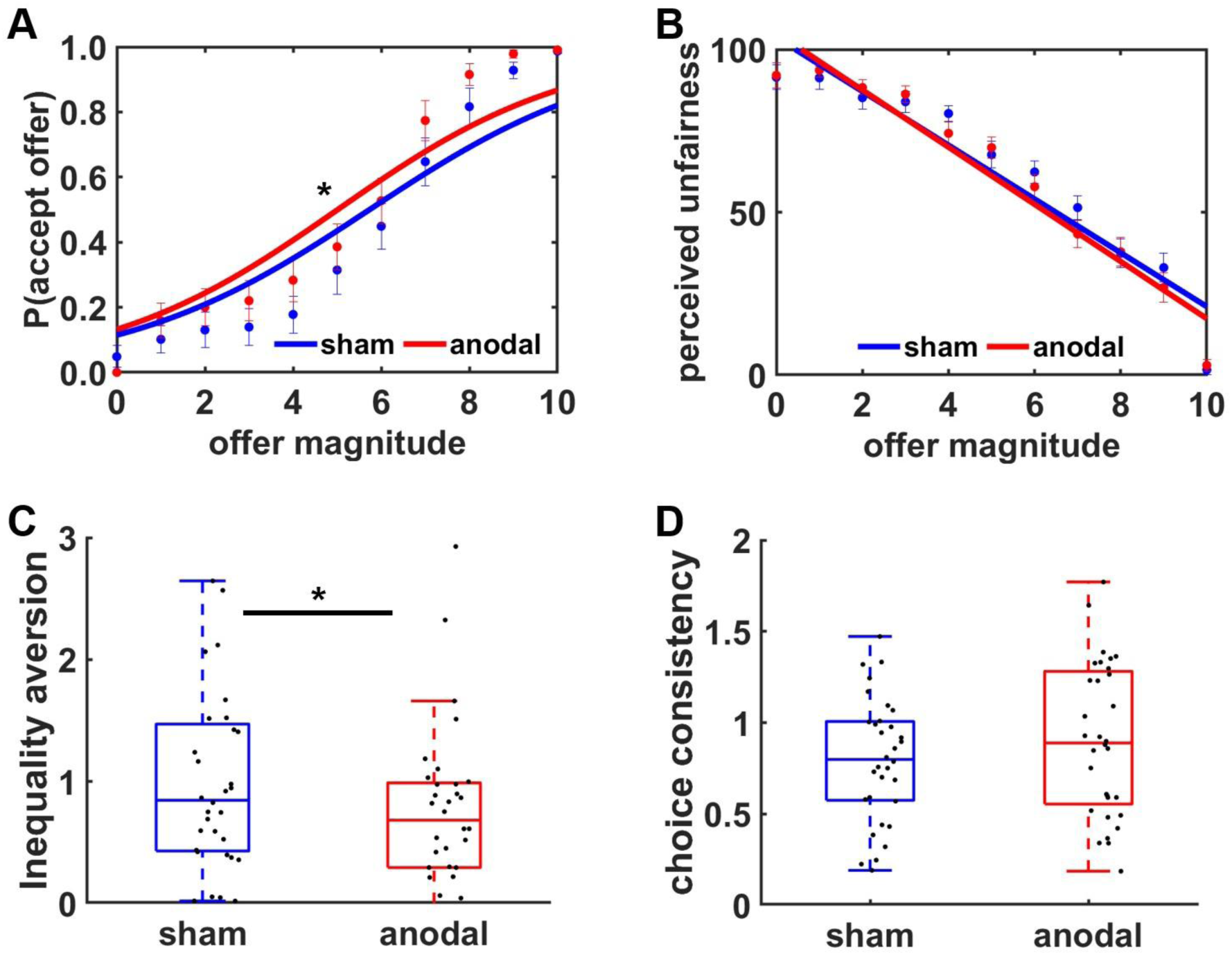
(A) Anodal versus sham tDCS over vmPFC increased acceptance rates independently of the magnitude of the monetary offer. (B) Higher offers were perceived as less unfair than lower offers, but there was no evidence that vmPFC tDCS changed the self-reported unfairness of offers. (C, D) Consistent with the observed higher acceptance rates under anodal tDCS, a computational model of inequality aversion suggests that increasing vmPFC excitability (C) significantly reduced aversion to disadvantageous inequality while leaving (D) choice consistency unaltered. Boxplots indicate inter-quartile ranges; black dots represent individual data points. Asterisks indicate significant effects (* *p* < 0.05).

To corroborate these findings, we fitted a computational model of inequality aversion to the choice data that quantified the aversion against disadvantageous unequal splits. Participants showed disadvantageous inequality aversion under sham, mean HDI = 0.41, 95%HDI = [0.02, 0.98], which was significantly reduced under vmPFC tDCS, mean HDI = - 0.27, 95%HDI = [-0.50, -0.06] (Figure 2C). In contrast, there were no stimulation effects on choice consistency, mean HDI = 0.11, 95%HDI = [-0.06, 0.32] (Figure 2D). This is relevant because previous studies related vmPFC activation to value representations and choice consistency (Bartra, McGuire, & Kable, 2013; Bowren, Croft, Reber, & Tranel, 2018), and strengthening such value representations through anodal tDCS should have resulted in increased choice consistency. Thus, converging model-based and model-free findings suggest a causal role of vmPFC for strengthening selfish interests over fairness concerns in normative choice.

We considered two alternatives that might explain the impact of vmPFC tDCS on inequality aversion: First, the vmPFC might directly be involved in down-regulating the emotional response to unfairness. According to the second possibility, the vmPFC does not affect the evaluation of unfairness per se but attenuates the influence of perceived unfairness on the decision process. These possibilities make dissociable predictions for the impact of tDCS on heart rate responses to unfairness as indicator of the reaction of the autonomous system to perceived unfairness: The first, emotion regulation-based alternative predicts that enhanced vmPFC excitability should reduce increases in heart rate in response to unfair offers. We first aimed to validate that changes in heart rate are indeed sensitive to unfairness: while the heart rate did not correlate with the objective amounts offered to participants or with the decision made, all *t* < 1.50, all *p* > 0.15, a separate GLMM suggested that heart rate changes were related to the self-reported perceived unfairness of offers, beta = 0.77, *t*(51) = 2.28, *p* = 0.03. This suggests that heart rates reflect the subjective unfairness of offers rather than monetary self-interests (which are related to the objective magnitude of an offer). Moreover, this physiological response to unfairness was significantly stronger for rejected compared with accepted offers, Unfairness × Choice: beta = -0.91, *t*(77) = 2.13, *p* = 0.04. Post-hoc tests suggest that perceived unfairness correlated with heart rate responses only for rejected, beta = 0.64, *t*(11) = 3.44, *p* = 0.006 (Figure 3A), not for accepted offers, beta = -0.16, *t*(21) = 0.77, *p* = 0.45 (Figure 3B). This is consistent with the interpretation that an increase in heart rate reflects the physiological response to offers perceived as unfair, which biases choices towards offer rejection. However, there was no significant influence of vmPFC tDCS on changes in heart rate, all *t* < 1.65, all *p* > 0.11. This does not support the first alternative according to which the vmPFC is involved in down-regulating physiological arousal in response to unfairness.

**Figure 3.**
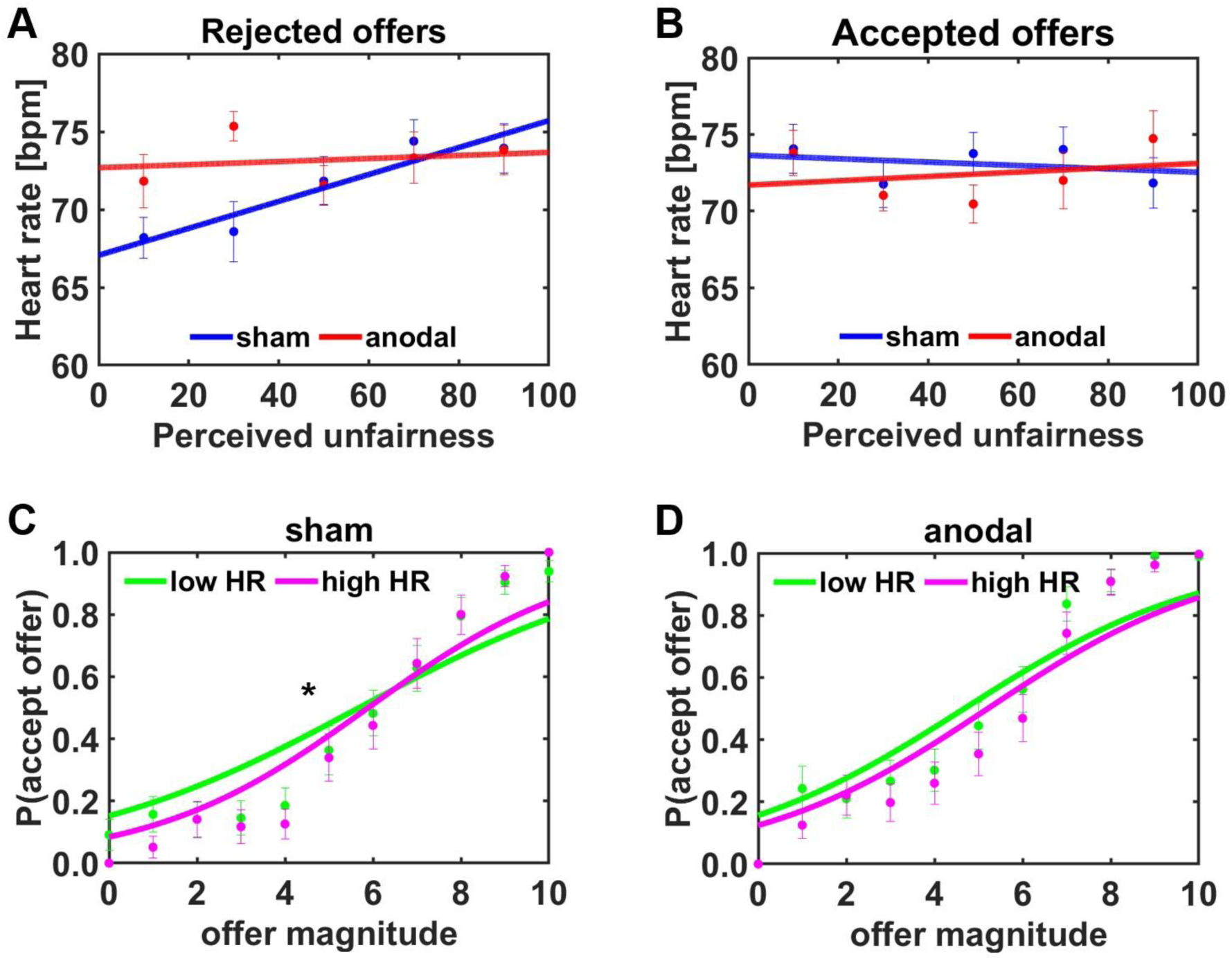
(A, B) Stimulation effects on heart rate changes during decision making. Under sham, heart rates significantly increased as a function of perceived unfairness for (A) rejected offers, but not for (B) accepted offers. We observed no significant effects of vmPFC tDCS on heart rates. Instead, vmPFC tDCS attenuated the influence of heart rates on decision making: (C) Under sham, higher hear rates during low, unfair offers increased the likelihood of rejecting the offer. (D) This influence of heart rates on decision making was reduced (and, in fact, no longer significantly different from zero) under anodal vmPFC tDCS.

According to the second alternative, vmPFC does not reduce the perceived unfairness of offers or the physiological arousal to unfairness but moderates how the physiological arousal influences choice behavior. To assess whether the impact of vmPFC tDCS on choices was moderated by heart rate changes, we added trial-by-trial measures of heart rate as further predictor to the model regressing binary choice on tDCS and Offer. Consistent with the findings reported above, increases in heart rates were associated with an enhanced likelihood of rejecting unfair offers, Offer × Heart rate, beta = 2.07, *z* = 4.33, *p* < 0.001. Importantly, this association between heart rate and choice behavior was reduced under vmPFC tDCS, tDCS × Offer × Heart rate, beta = -1.32, *z* = 2.36, *p* = 0.02. Separate GLMMs for sham and anodal tDCS suggest that faster heart rates were linked to higher rejection rates for low, unfair offers only under sham, beta = 1.83, *z* = 3.96, *p* < 0.001 (Figure 3C), not under anodal tDCS, beta = 0.57, *z* = 1.24, *p* = 0.22 (Figure 3D). This suggests the vmPFC to play a causal role in attenuating the relationship between physiological responses to unfairness and decisions to reject the offer. To corroborate these findings, we tested whether vmPFC stimulation moderates the influence of physiological arousal also in our computational model of inequality aversion by adding heart rate as covariate to the computational model described above. In fact, under anodal tDCS the utility of accepting unfair offers was increased for higher heart rates, mean HDI = 0.04, 95%HDI = [0.01, 0.09]. Taken together, this provides converging evidence that the vmPFC is causally involved in promoting the acceptance of unfair offers associated with high physiological arousal.

### vmPFC tDCS increases decision confidence after self-benefitting choices

Given that anodal vmPFC tDCS reduced the influence of unfairness-related physiological arousal on decision making, we next asked whether the vmPFC affects not only choices themselves but also the post-decision confidence in having made the best choice. Under sham, post-decision confidence was higher for rejected compared with accepted offers, beta = -18.03, *t*(26) = 2.12, *p* = 0.04. Confidence was also increased the less money was offered to participants, beta = -12.51, *t*(24) = 6.62, *p* < 0.001, particularly if the unfair offer was rejected rather than accepted, Offer × Choice: beta = 40.28, *t*(28) = 7.12, *p* < 0.001. Participants were therefore less confident to have made the optimal choice after accepting versus rejecting unfair offers. This effect was significantly reduced after vmPFC tDCS, tDCS × Offer × Choice: beta = -12.21, *t*(26) = 2.74, *p* = 0.01 (Figure 4). Separate GLMMs for accepted and rejected offers suggest that vmPFC tDCS increased confidence after offer acceptance, tDCS: beta = 6.66, *t*(23) = 2.17, *p* = 0.04, particularly for unfair offers, tDCS × Offer: beta = -8.79, *t*(25) = 2.66, *p* = 0.01. In contrast, there was no evidence for tDCS effects on confidence after offer rejections, all *t* < 0.79, all *p* > 0.45. Taken together, enhancing vmPFC excitability increased both the willingness to accept unfair offers and the subjective post-decision confidence that accepting an offer was the best choice.

**Figure 4.**
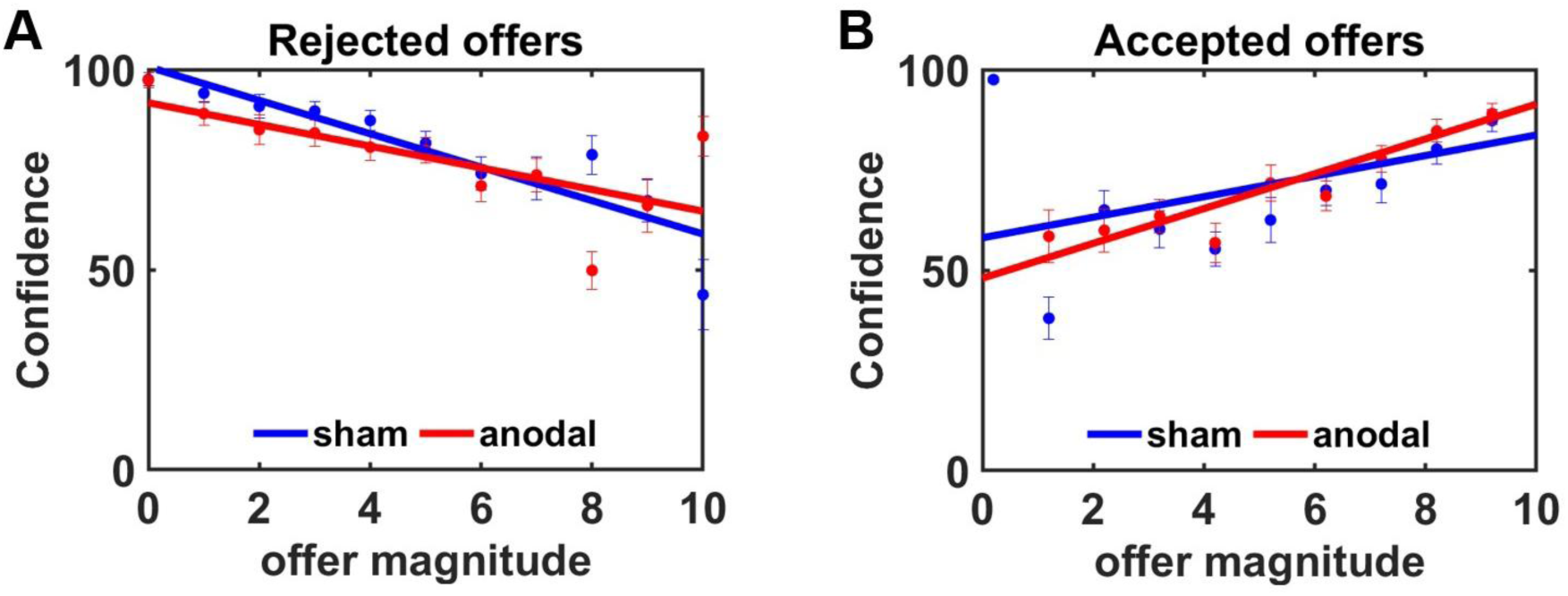
Influence of vmPFC tDCS on post-decision confidence. (A) Confidence was higher after rejected offers the smaller the magnitude of the offered amount. There was no evidence for stimulation effects on confidence after rejected offers. (B) In contrast, confidence after offer acceptance was higher for increasing monetary offers, and the influence of offer magnitude on confidence was significantly enhanced under vmPFC tDCS. This suggests vmPFC tDCS to increase confidence for large, accepted offers.

## Discussion

The vmPFC was ascribed a central role in integrating emotions into the decision process, but it remained unknown whether vmPFC influences decisions via directly down-regulating emotion-related physiological arousal or via attenuating the impact of the physiological arousal on the decision process. Here, we provide evidence that the vmPFC is causally involved in attenuating the influence of unfairness-related physiological arousal on decisions involving trade-offs between selfish interests and fairness considerations. First, we found the self-reported perceived unfairness of rejected offers, but not the monetary self-interest associated with offers, to be linked to increased heart rates response, consistent with the idea that changes in heart rate reflect the physiological response of the autonomous nervous system to unfairness (Dunn et al., 2012). There was no evidence for an influence of vmPFC tDCS on the perceived unfairness of offers or on bodily arousal, which does not support the view that vmPFC directly affects normative evaluations of unfairness or inhibits physiological arousal related to unfairness. Instead, our findings suggest that enhancing vmPFC excitability attenuates the influence of unfairness-related heart rate changes on rejection decisions in the Ultimatum game. Higher heart rates biased the rejection of unfair offers only under sham, whereas under anodal tDCS changes in heart rate showed no significant influence on normative decisions. While previous neuroimaging and lesion research yielded inconclusive results regarding the vmPFC’s role for trading-offs between fairness and self-interest (Baumgartner et al., 2011; Feng et al., 2015; Gilam et al., 2017; Gu et al., 2015; Koenigs & Tranel, 2007; Lo Gerfo et al., 2019), the current high-definition stimulation protocol provides evidence that vmPFC promotes the maximization of one’s self-interest by lowering the sensitivity to fairness violations relative to selfish interests. The spatial precision of our high-definition tDCS setup that minimizes stimulation effects on regions such as DLPFC is a strength compared to previous vmPFC tDCS studies, though we note that we cannot fully rule out tDCS effects on lateral orbitofrontal cortex or subgenual anterior cingulate cortex. In any case, the current results are consistent with findings from other domains of decision making according to which vmPFC reduces the influence of action costs like risk, losses, or waiting time on decision making (Cecchi et al., 2024; Ciaramelli et al., 2021; Kroker et al., 2022), suggesting that vmPFC may generally strengthen reward maximization over cost avoidance.

The vmPFC may be able to moderate the influence of emotional arousal on decision making due to its central role in a distributed neural decision network. The vmPFC receives input from several regions involved in value computation, including insula, dlPFC, and striatum (Ghaziri et al., 2017; Jackson, Bajada, Lambon Ralph, & Cloutman, 2020). The insula was related to emotion processing (Gogolla, 2017), including violations of social norms (Harle, Chang, van ’t Wout, & Sanfey, 2012; Sanfey, Rilling, Aronson, Nystrom, & Cohen, 2003). In addition, insula activation and connectivity between insula and vmPFC play a role for the interoceptive awareness of bodily signals (Craig, 2002; Critchley & Harrison, 2013; Jarrahi et al., 2015; Kirk, Gu, Harvey, Fonagy, & Montague, 2014; Sagliano et al., 2019). Interoceptive awareness of unfairness-related bodily arousal was shown to reduce the influence of unfairness on social decisions (Kirk et al., 2011), indicating that interceptive awareness of physiological arousal may allow decoupling emotional responses from the decision process. This suggests that vmPFC tDCS might have reduced inequality aversion via enhancing introspection, thereby reducing the impact of unfairness-elicited physiological arousal on decision making. We note that fairness considerations may influence social decisions not only via unfairness-triggered emotional responses; for example, the right dlPFC was hypothesized to encode abstract representations of social norms (Knoch, Pascual-Leone, Meyer, Treyer, & Fehr, 2006), and disrupting dlPFC activation reduces the value assigned to punishing norm violations in vmPFC (Baumgartner et al., 2011). Moreover, other studies observed vmPFC stimulation to down-regulate emotion-related physiological arousal in tasks where participants were explicitly instructed to direct attention to their emotions (Hilz et al., 2006; Szeska, Punjer, Riemann, Meinzer, & Hamm, 2022). In situations requiring no intentional down-regulation of emotions such as in value-based decision making, vmPFC may shield value computations from intrusion by decoupling physiological arousal from the decision process rather than by directly inhibiting these bodily signals.

Besides value computations, the vmPFC was also related to representing confidence judgements (De Martino, Fleming, Garrett, & Dolan, 2013; Shapiro & Grafton, 2020; Vaccaro & Fleming, 2018). Our data show for the first time that vmPFC is in fact causally involved in computations of confidence, as anodal vmPFC tDCS increased decision confidence for accepted unfair offers. Although it was suggested that separate neural signals in vmPFC encode value and confidence (Lebreton, Abitbol, Daunizeau, & Pessiglione, 2015), the confidence in having made the best choice scales with the difference in value between choice options (Bobadilla-Suarez, Guest, & Love, 2020; Shapiro & Grafton, 2020). By attenuating the influence of arousal on decision making, vmPFC tDCS may have increased both the value of accepting unfair offers (as suggested by our computational model of inequality aversion) and subjective post-decision confidence after accepting unfair offers.

Taken together, the current findings provide a mechanistic understanding of the vmPFC’s contribution to computing trade-offs between social norms and self-interests. The vmPFC is causally involved in decoupling physiological arousal triggered by fairness violations from the decision process, thereby reducing aversion to disadvantageous inequality and strengthening subjective confidence when accepting unfair offers. This advances our knowledge of the brain mechanisms causally underlying the interaction between emotions and cognition in social and potentially also non-social domains of decision making.

## Acknowledgements

AS received support from an Emmy Noether fellowship from the German Research Foundation (SO 1636/2-2) and an Exploration grant from the Boehringer Ingelheim Foundation. All authors declare to have no conflicts of interest.

## Data availability statement

The raw data underlying the findings are available on Open Science Framework (OSF; https://osf.io/d6m54/overview?view_only=36364c95eab747f6b4e6292d6cab3be6).

